# Entrainment of Endangered Sturgeon by a Large Water Diversion: Rescue, Enumeration, and Conservation Opportunities

**DOI:** 10.1101/2021.07.22.453439

**Authors:** K. Jack Killgore, Jan Jeffrey Hoover, William Todd Slack, Steven G. George, Christopher G. Brantley

**Affiliations:** U. S. Army Corps of Engineers, Engineer Research and Development Center, Environmental Laboratory, Vicksburg, Mississippi, United States of America; U. S. Army Corps of Engineers, New Orleans District, New Orleans, Louisiana, United States of America

## Abstract

The Bonnet Carre’ Spillway diverts water from the Mississippi River through a floodway into Lake Pontchartrain to reduce river stages at New Orleans and prevent flood damages. Pallid Sturgeon, a federally listed species under the Endangered Species Act, and Shovelnose Sturgeon, listed under the Similarity of Appearance rule, are entrained through the Spillway structure and become trapped in the Spillway canals and other waterbodies. Five openings and corresponding rescue operations occurred between 2008 and 2019 after each Spillway closure. Operational parameters spanned a range of water temperatures and seasons with magnitude and duration of discharge varying across all openings. A total of 70 days with crew number ranging from 6 to 12 were expended to rescue 57 Pallid Sturgeon and 362 Shovelnose Sturgeon after the five openings that spanned 240 total days. More sturgeon were entrained at higher water temperatures, with greater numbers of bays opened, and for longer periods of time. Recovery of sturgeon is initially high but over time declines as sturgeon are depleted from the floodway, stranded in isolated waterbodies in the floodway, and/or displaced further downstream into Lake Pontchartrain during longer openings. Sturgeon that cannot find their way back to the floodway are unlikely to be rescued. Recent population studies indicate that less than 1% of the total population size in the Lower Mississippi River are entrained. However, this does not take into account those individuals entrained but not captured and the potential impacts of more frequent openings of the structure. Conservation recommendations are provided to increase catch efficiency and recovery of the endangered sturgeon.

## Introduction

Large-scale interbasin water transfer projects occur worldwide for various purposes including domestic water supply, energy production, agricultural irrigation, marsh restoration, and flood control (Sternberg 2016; Shumiloval 2018). One of the largest interbasin freshwater diversions in the United States is the Bonnet Carré Spillway (BCS) on the Lower Mississippi River 53 river kilometers upstream of New Orleans, LA (Figure 1). The BCS structure, constructed by the U. S. Army Corps of Engineers as a flood risk management feature following the Mississippi River Flood of 1927, is a needle-controlled dam 2347 m in length and the design discharge capacity is 7079 cubic meters per second (cms) (U.S. Army Corps of Engineers 1998). It diverts water from the Mississippi River to prevent river discharges downstream towards New Orleans in excess of 35,400 cms, into a floodway that empties into Lake Pontchartrain, a shallow, brackish lagoon. Typically, Mississippi River water is < 2 ppt salinity (USGS Gage 7374000, Mississippi River at Baton Rouge), whereas in Lake Pontchartrain ranges from 1.2 to 5.4 ppt salinity (Sikora and Kjerfve 1985).

**Figure 1.**
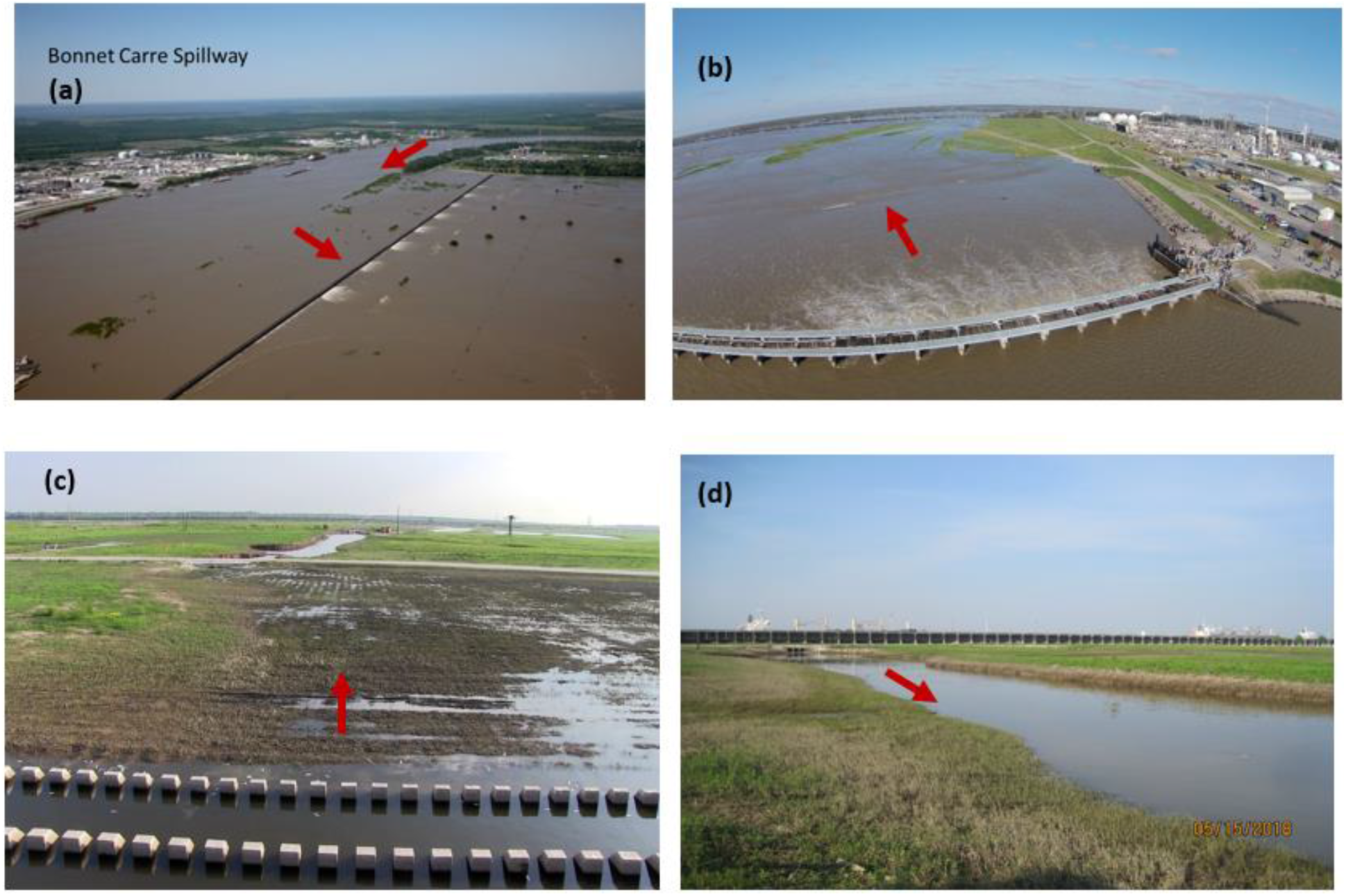
Bonnet Carré Spillway structure, stilling basin, and outflows. Red arrows indicate direction of flow: aerial looking north upriver and through the structure during the 2011 opening (a), aerial looking east from structure towards Lake Pontchartrain during the 2011 opening (b), ground looking east from stilling basin towards Lake Pontchartrain after closure (c), and ground looking west at Barbar’s Canal toward structure after closure (d). Aerial views provided by USACE New Orleans District.

As of 2019, the BCS has been operated fourteen times (Figure 2). Frequency of operations between 1937 and 2008 occurred at 2-23 year intervals, most between 4-14 years, and averaged overall once every 8.9 years, or just over 1% during that time period. Duration of openings ranged from 13 days in 1975 to 79 days during the second opening of 2019. Number of bays open each day ranged from 1935 in 1975 to 22204 in 1973. Since 2008, operations have occurred at 1-5 year intervals, averaging 2.7 years overall, or about 5% during that time period. Record floods have occurred in the Lower Mississippi River over the last decade necessitating openings of the BCS at a three-five times greater frequency than historic operations. In addition to increased frequency of operation, magnitude and duration of flooding are also increasing. The 2011 flood set new stage records at multiple locations along the Lower Mississippi River. The 2019 flood was the longest in modern history, and for the first time, the Spillway was opened twice in the same year.

**Figure 2.**
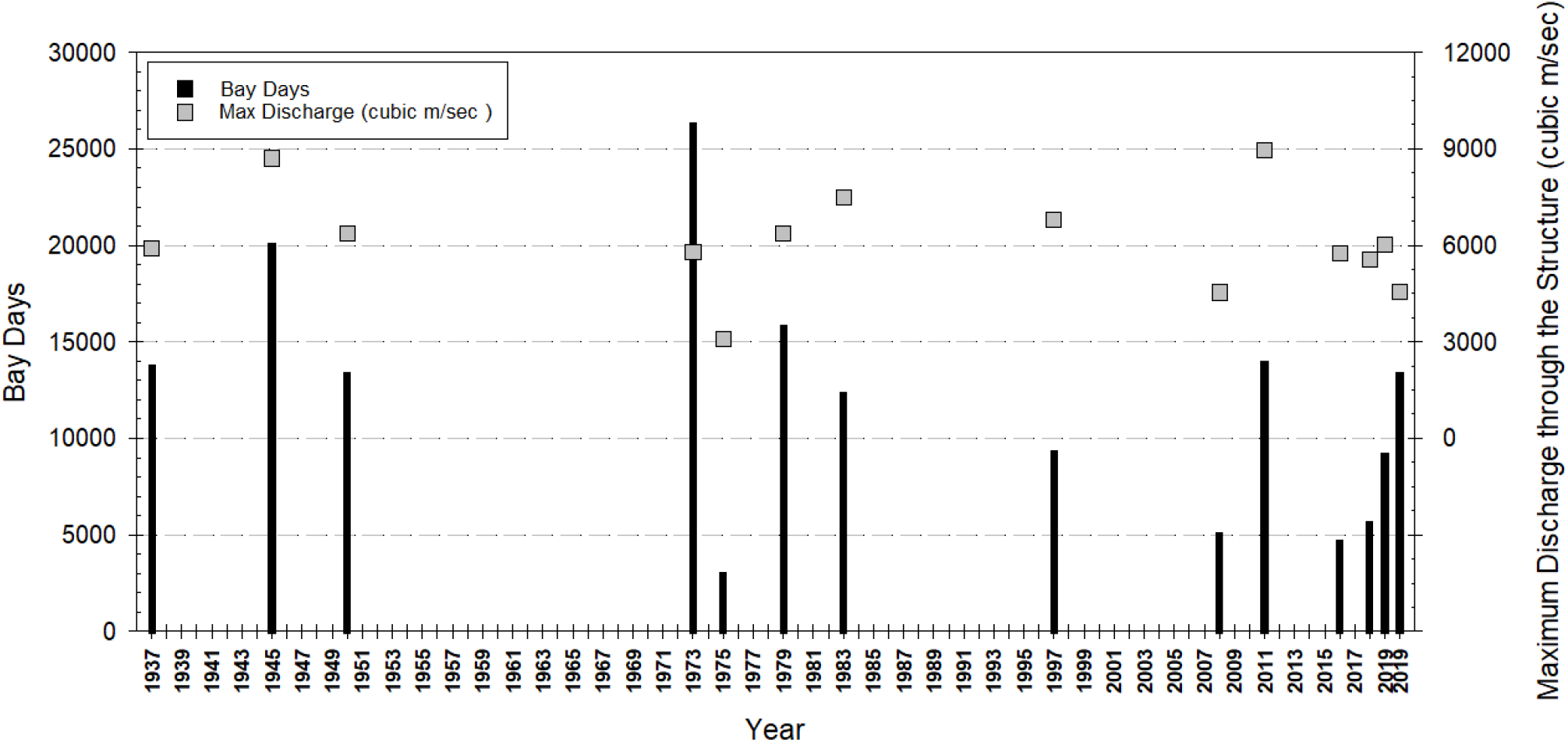
Frequency, maximum discharge, and bay days (i.e., number of bays open per day) for the 14 openings of the Bonnet Carré Spillway since the structure was constructed in 1931. Sturgeon sampling occurred after the structure was closed from 2008 to 2019. The structure was open twice during 2019.

Entrainment risk has been evaluated for Green Sturgeon (*Acipenser medirostris*) through agricultural diversion pipes (Mussen et al 2014), Lake Sturgeon through a hydroelectric station (McDougall et al. 2014), White (*Acipenser transmontanus*) and Pallid Sturgeon (*Scaphirhynchus albus*) in the vicinity of dredges (Boysen and Hoover 2009; Hoover et al. 2011), and an overall synthesis on interactions between sturgeon and water resource development including sturgeon entrainment and impingement (Cooke et al. 2020). However, effects of flood control diversion openings on imperiled sturgeon species in the Mississippi River have not been described. The fish cannot pass back into the river once water level drops sufficiently below the backside of the weir sill. This can happen as the BCS structure is closed, restricting water into the floodway or with bays open on the structure and falling stages on the Mississippi River. It is believed that as a lotic freshwater species, sturgeon will be unable to survive long-term in the lentic brackish environments of the Lake Pontchartrain system. Therefore, it is assumed conservatively that entrained sturgeon represent impacts to the source population in the river.

After the structure was closed in 2008, a decision was made by the U.S. Army Corps of Engineers, New Orleans District to sample water bodies within the BCS for Pallid Sturgeon (*Scaphirhynchus albus*), listed federally as endangered under the Endangered Species Act, as well as the Shovelnose Sturgeon (*Scaphirhynchus platorynchus*), which are sympatric with Pallid Sturgeon and in 2010 were listed as a threatened species under the “Similarity of Appearances” provisions of the Endangered Species Act only in those areas where they co-occur with Pallid Sturgeon (Figure 3). Under section 7 of the Endangered Species Act, all federal agencies are responsible to ensure that actions they authorize, fund, or carry out are not likely to jeopardize the continued existence of any listed species. Principal objectives of this effort were to rescue entrained Pallid Sturgeon, return them to the Mississippi River, and to enumerate impacts to the population. Rescue operations comply with the ESA and have been found to be effective for other species of sturgeon. Telemetry and modeling studies showed that rescue can provide nearly complete mitigation for population impacts of entrained Green Sturgeon (Thomas et al. 2013).

**Figure 3.**
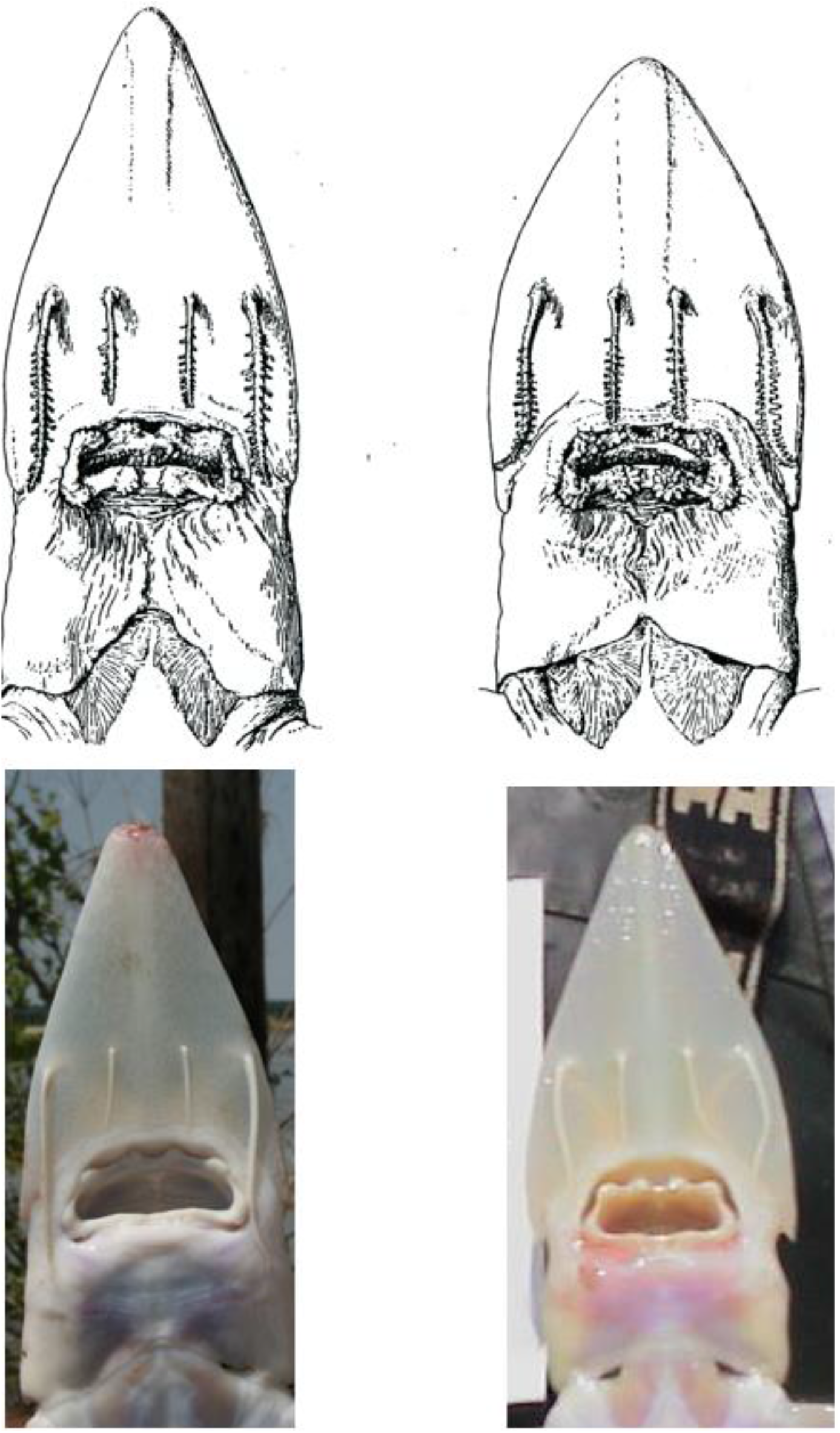
Ventral view of Pallid (left) and Shovelnose (right) Sturgeon of the Lower Mississippi River. Pallid Sturgeon have shorter inner barbels relative to outer barbels, longer head, and larger mouth. Top drawings are from Forbes and Richardson (1905).

Within the first hour of sampling in 2008, a Pallid Sturgeon was captured in a BCS canal downstream of the structure inaugurating a series of sampling and rescue events after the structure was operated each time in 2008, 2011, 2016, 2018, and 2019. Intense and equitable sampling was required during each operation to evaluate impacts to the species and to evaluate the relationships between numbers of sturgeon entrained with functional parameters of the structure, specifically onset, magnitude, and duration of flooding and operation. This article summarizes the number of sturgeon collected after each of the five openings. It also provides data on sturgeon movements and habitat quality. It identifies differences in catch among the openings based on environmental and operational conditions, estimates potential impacts to population abundance, and recommends sampling requirements and strategies for future openings to maximize catch and minimize mortality of entrained sturgeon.

## Study Area

The BCS was built in 1929-31 by the U.S. Army Corps of Engineers on the east side of the Mississippi River near the site of the former Bonnet Carré Crevasse 53 river km above New Orleans. It is recognized by the American Society of Civil Engineers as a Historic Civil Engineering Landmark and eligible for listing on the National Register of Historic Places. The BCS structure is 2347 m long with 350 concrete bays (or weirs) each 6.1 m in length. There are 176 bays with a weir elevation of 5.15 m N.G.V.D (National Geodetic Vertical Datum) (i.e., high bays) and the remaining 174 bays have a weir crest elevation of 4.54 m N.G.V.D. (i.e., low bays). Each bay is closed with 20 timber needles (also referred to as pins) measuring 29-30 cm in width and either 3 m or 3.7 m in length depending on the elevation of the weir crest. Needles are inserted vertically across each bay while the Spillway is closed. During openings, a travelling gantry crane mounted on narrow-gage tracks on top of the BCS structure lift the pins from the bay allowing Mississippi River water to pass unimpeded into the floodway (Figure 1). Pins are stored above each bay during operations. Discharge through the BCS is regulated by the number of bays opened or closed on the structure. The bays allow flow directly into a stilling basin approximately 15 m wide with three rows of low concrete baffles next to a heavy articulated concrete mat 53 to 69 m wide to dissipate the flow energy. Floodwaters are then directed into a leveed floodway conveying water from the weir structure into Lake Pontchartrain.

The 3085 hectare floodway is 9.2 km in length confined by levees that are 2.3 km wide at the river end and 3.8 km wide at the lake end. According to the Master Plan (USACE 1998), the lands in the floodway are characteristic of an alluvial floodplain that vary in elevation from 3 meters near the river to mean sea level at Lake Pontchartrain. The Convent-Commerce soils series consist of soft organic clays with layers of silt and peat, and high water content that support grasses and sedges. Bottomland hardwood forest comprise approximately 40% of the total project acreage while the remainder of the floodway is mostly disturbed land with little vegetation following opening and closing of the structure. When the structure is closed but still leaking through the pins, the Y-canal and Barbar’s canal drain the majority of water into two large borrow canals at the lower half of the floodway that empty into Lake Pontchartrain (Figure 4). In addition, over 25 shallow ponds created by sand excavation activities are scattered across the floodway.

**Figure 4.**
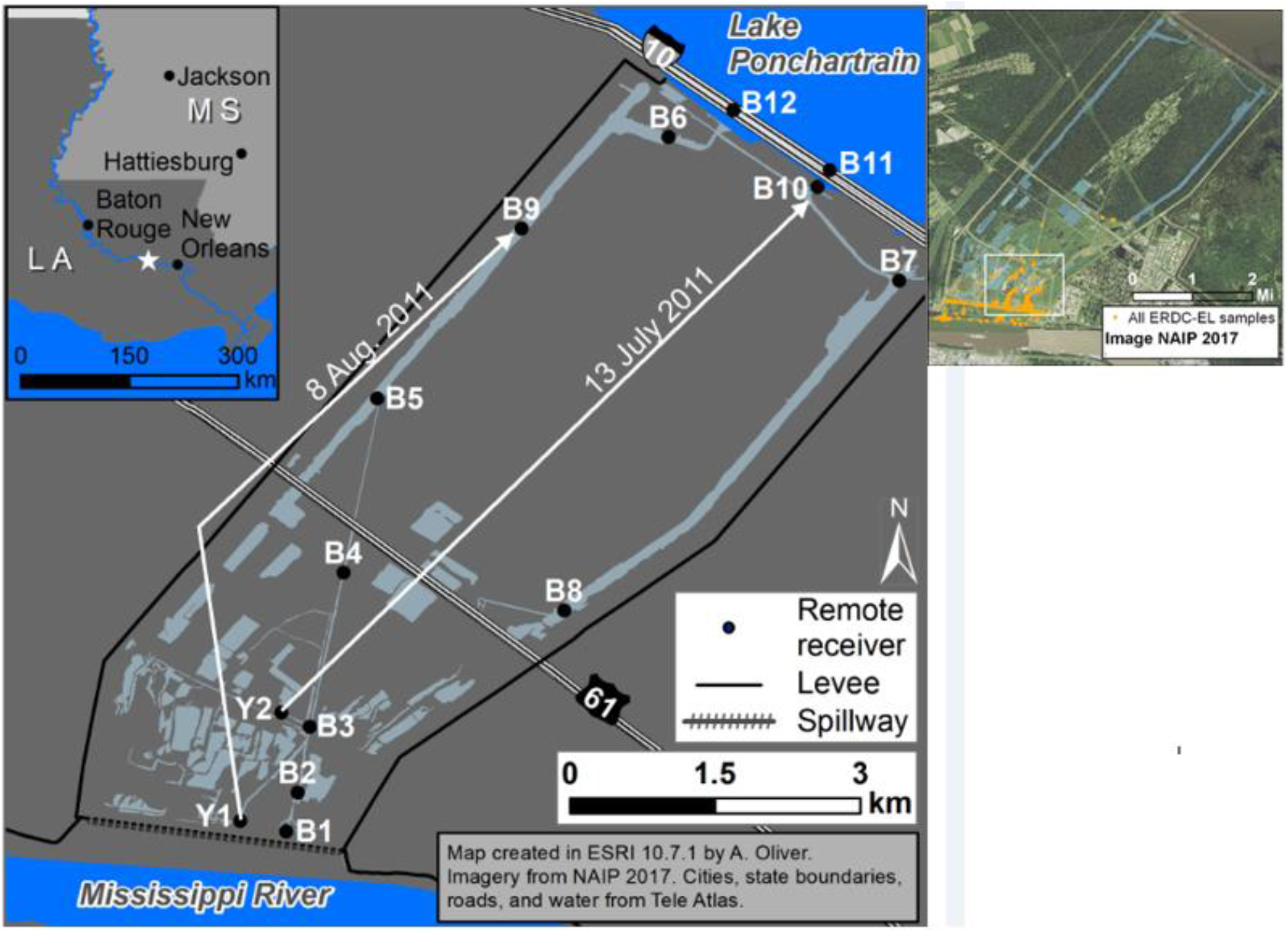
Sampling locations in the Bonnet Carré Spillway. The smaller inset map displays the concentration of sturgeon sampling during all openings from 2008 - 2019. The larger map displays the location of acoustic telemetry receivers in 2011 and arrows indicated those receivers moved to new locations. The letter “Y” refers to the Y-Canal and the letter “B” refers to Barbar’s canal and receiving canals. The blue color denotes all waterbodies in the floodway after closure of the structure. The stilling basin is immediately below the Spillway structure.

The structure is operated once the discharge on the Mississippi River reaches and is expected to exceed 35400 cms. After structure closure, if Mississippi River levels are above the concrete weir, water can continue to pass over the weir and through the openings between pins in the closed bays. This movement of water through the closed structure is commonly called leakage and can often maintain a shallow (0.5 – 1 m) sheet flow across upper portions of the floodway. Pin leakage at high elevation bays cease flowing at 5.15 m at the Bonnet Carre’ gage, which also eliminates sheet flow and allows vehicular access to sampling sites. Low bays continue to leak until gage reading is at 4.54 m eliminating flows through the various canals and ditches. The period of leakage varied each year and was an important consideration for sampling strategies.

## Materials and Methods

The floodway was sampled after five BCS operations: 2008, 2011, 2016, 2018 and 2019. Each operation was different in duration, magnitude of discharge passing through the structure, and time of year (Table 1). There were two separate operations in 2019 resulting in a total of 123 open days. The 2019 Mississippi River flood was the longest in recorded history reflected by the duration of the two openings. The highest maximum discharge through the structure occurred in 1945 with a discharge of 9005 cms followed by a discharge of 8946 cms during the 2011 opening that lasted 43 days. The 2011 flood set new records on maximum discharge at several gages in the Lower Mississippi River resulting in the higher discharge passed through the Spillway. The remaining three years had shorter openings (<30 days) and reduced maximum discharge (<6100 cms).

**Table 1.**
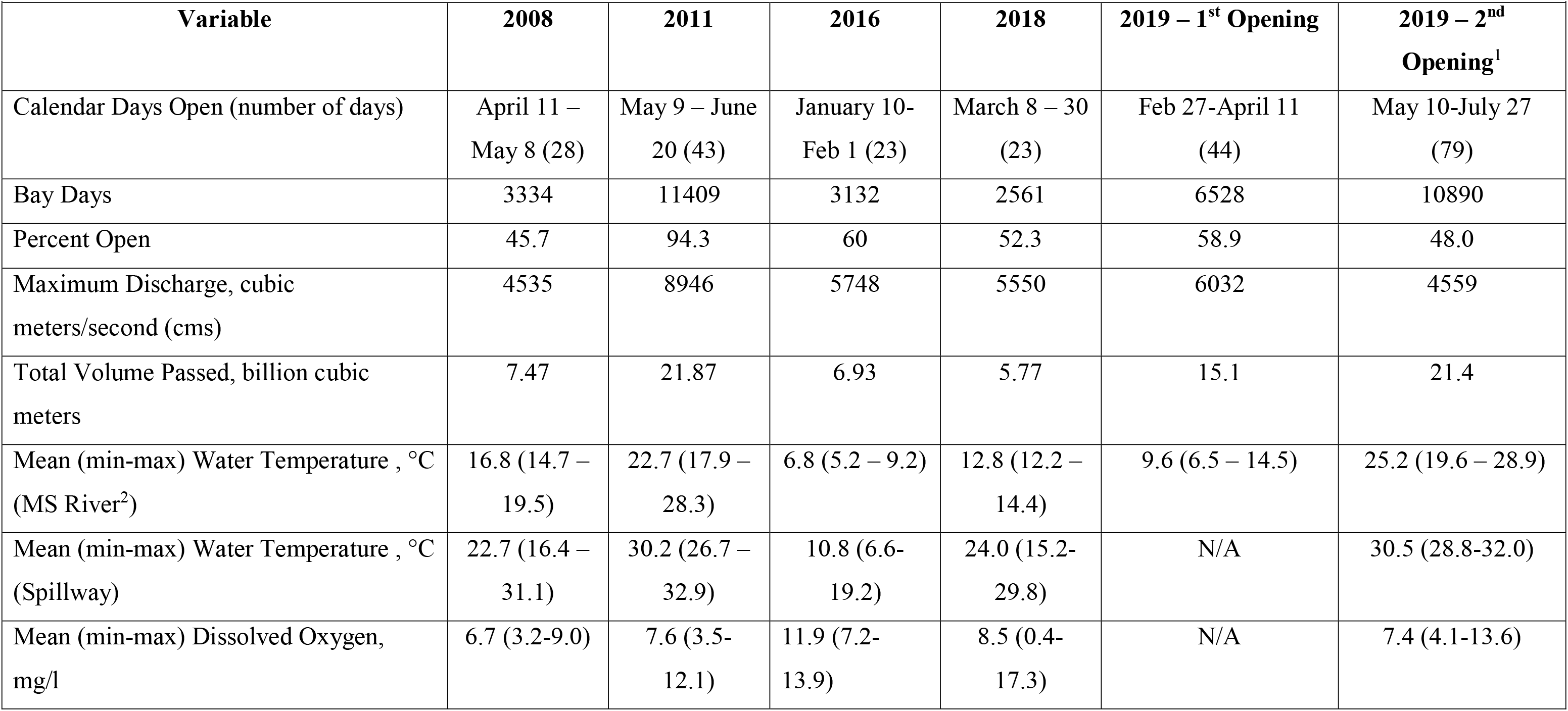
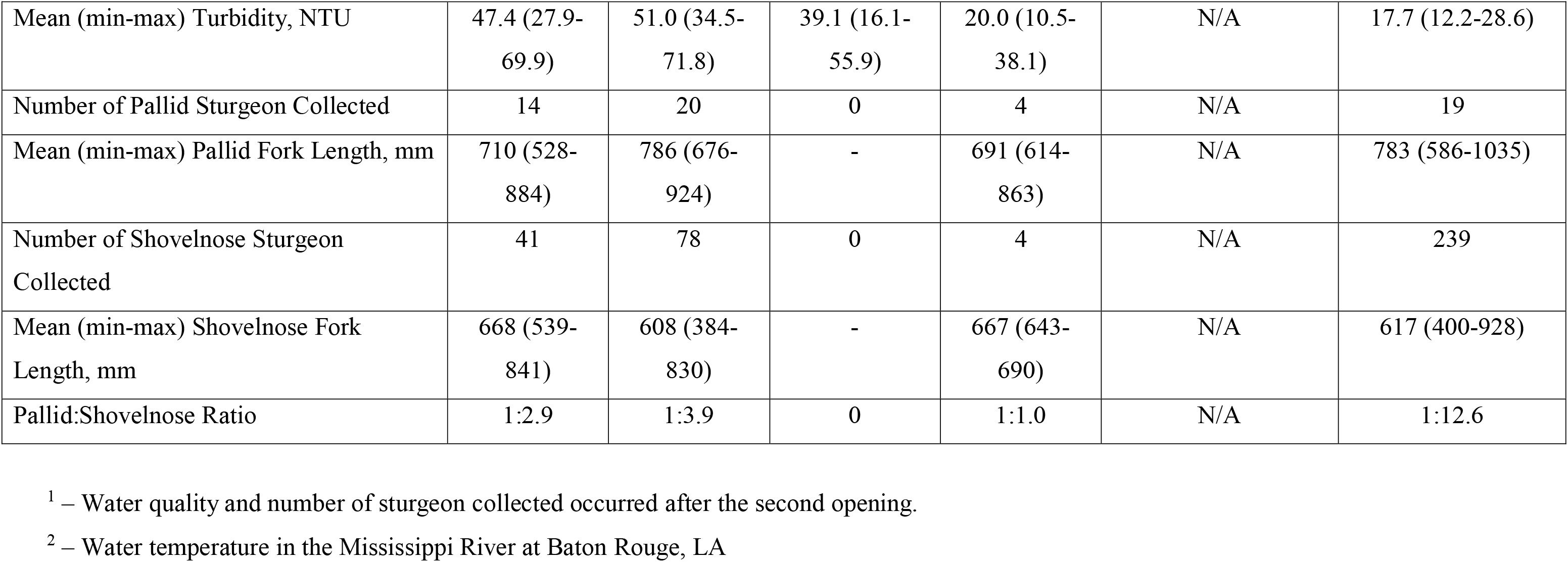
Characteristics of the Bonnet Carré Spillway after closure from 2008 – 2019 and number of sturgeon collected. Water quality measurements occurred after closure during sampling and included canals, ditches, and lakes.

| **Variable** | **2008** | **2011** | **2016** | **2018** | **2019 – 1<sup>st</sup> Opening** | **2019 – 2<sup>nd</sup> Opening<sup>1</sup>** |
| --- | --- | --- | --- | --- | --- | --- |
| Calendar Days Open (number of days) | April 11 – May 8 (28) | May 9 – June 20 (43) | January 10- Feb 1 (23) | March 8 – 30 (23) | Feb 27-April 11 (44) | May 10-July 27 (79) |
| Bay Days | 3334 | 11409 | 3132 | 2561 | 6528 | 10890 |
| Percent Open | 45.7 | 94.3 | 60 | 52.3 | 58.9 | 48.0 |
| Maximum Discharge, cubic meters/second (cms) | 4535 | 8946 | 5748 | 5550 | 6032 | 4559 |
| Total Volume Passed, billion cubic meters | 7.47 | 21.87 | 6.93 | 5.77 | 15.1 | 21.4 |
| Mean (min-max) Water Temperature , °C (MS River<sup>2</sup>) | 16.8 (14.7 – 19.5) | 22.7 (17.9 – 28.3) | 6.8 (5.2 – 9.2) | 12.8 (12.2 – 14.4) | 9.6 (6.5 – 14.5) | 25.2 (19.6 – 28.9) |
| Mean (min-max) Water Temperature , °C (Spillway) | 22.7 (16.4 – 31.1) | 30.2 (26.7 – 32.9) | 10.8 (6.6- 19.2) | 24.0 (15.2- 29.8) | N/A | 30.5 (28.8-32.0) |
| Mean (min-max) Dissolved Oxygen, mg/l | 6.7 (3.2-9.0) | 7.6 (3.5- 12.1) | 11.9 (7.2- 13.9) | 8.5 (0.4- 17.3) | N/A | 7.4 (4.1-13.6) |
| Mean (min-max) Turbidity, NTU | 47.4 (27.9-69.9) | 51.0 (34.5-71.8) | 39.1 (16.1-55.9) | 20.0 (10.5-38.1) | N/A | 17.7 (12.2-28.6) |
| Number of Pallid Sturgeon Collected | 14 | 20 | 0 | 4 | N/A | 19 |
| Mean (min-max) Pallid Fork Length, mm | 710 (528-884) | 786 (676-924) | - | 691 (614-863) | N/A | 783 (586-1035) |
| Number of Shovelnose Sturgeon Collected | 41 | 78 | 0 | 4 | N/A | 239 |
| Mean (min-max) Shovelnose Fork Length, mm | 668 (539-841) | 608 (384-830) | - | 667 (643-690) | N/A | 617 (400-928) |
| Pallid:Shovelnose Ratio | 1:2.9 | 1:3.9 | 0 | 1:1.0 | N/A | 1:12.6 |

<sup>1</sup> – Water quality and number of sturgeon collected occurred after the second opening.
<sup>2</sup> – Water temperature in the Mississippi River at Baton Rouge, LA

### Collecting Techniques

The goal was to capture as many Pallid Sturgeon as possible in the BCS and release them back into the Mississippi River. Collecting effort occurred primarily in the upper end of the BCS including the canals (Barbar’s and Y), ditches, and stilling basin once the high bays ceased leaking (Figure 4). Floodway lakes were also periodically sampled. Multiple gears and techniques were used in the various waterbodies and collecting efforts ceased 1-2 days after the last sturgeon was collected. Collecting efforts were repeated the following week(s) if flow persisted in the canals due to bay leakage.

1. Boat-mounted electroshocker – Operated with DC pulse at an output of 4-6 amps at 60 Hz (targets wide size range of fishes) or 1-2 amps at 15 Hz (targets larger benthic fishes) using a Smith-Root 7.5 GPP system. Two dip-netters on the bow, and often one behind them, would scan the water surface for immobilized fish, collect all sturgeon, and collect a representative sample of the fish assemblage. Sturgeon were visible at the surface for only a few seconds requiring quick reflexes by the dippers. Unit of effort was expressed as minutes electroshocked.
2. Gill Nets – Three types of gillnets were used and unit of effort for each was expressed as hours fished:

a. A 15.2 to 18-m section of a trammel net (40-m long, 2.4-m deep, with 6.3 cm square mesh) was set at the end of a 200 m reach of Barbars Canal between upstream road/culvert crossing and downstream mid-water pipeline crossing prior to electroshocking and used as a block net to capture or contain fishes during sampling.
b. Sets of 43 x 3 m experimental mesh (7.6 – 15.2 cm square mesh) gillnets were set in lakes prior to electroshocking.
c. A 30.5 x 1.8 m net with 7.6 – 10.2 inch square mesh was set in lakes prior to electroshocking.
3. Conventional seine - Smaller ditches were occasionally sampled with a conventional 6.1 m by 2.4-m seine of 1-cm bar mesh. Unit of effort was expressed as number of hauls
4. Modified gillnet-seine - A section of a 27.4 m X 1.8 m gill net was used to herd and capture fishes in the Stilling Basin. A block net with 7.6-cm square mesh was set in the Stilling Basin at some distance away from the seining operation to increase containment during seining. The gill net seine had square mesh size ranging from 1.9 to 6.4-cm tied to bamboo brails on each end. Larger mesh minimized entanglement of spines from small Blue Catfish (*Ictalurus furcatus*) that were abundantly distributed in the Stilling Basin. A crew would begin on one end of the Stilling Basin and pull the seine through the larger channel while other personnel in the two smaller channels between the two rows of baffles would splash the water to herd fish towards the seine or capture fish with large dipnets. The crew would periodically stop at the block net to remove and record fish. The block net was moved further away and seining commenced again until the entire Stilling Basin was sampled. Unit of effort was expressed as hours seined.
5. Visual Sightings – Sturgeon were visually sighted in the Stilling Basin by the ground team or from the gantry crane on top of the structure. Once sighted, the ground team waded into the basin with castnets or large dipnets to capture the individual. Dead sightings were also occasionally made in other waterbodies and sturgeon were retrieved.
6. Benthic trawls, hoop nets, and trotlines were also used in the flowing channels, but these gears were eventually dropped due to low catches attributed to entanglement on bottom obstructions and trash entangled in the nets and trotline hooks.

### Sturgeon Identification

Sturgeon were identified to species, enumerated, and fork length recorded. Discriminating between Pallid Sturgeon and Shovelnose Sturgeon can be challenging, however (Figure 3). They co-occur, are morphologically similar and known to hybridize, and they vary genetically throughout their range (Schrey et al. 2011; Jordan et al. 2019), but results of genetic studies and of morphological studies frequently yield inconsistent results (e.g., Schrey et al. 2007 vs Ray et al. 2007; Bailey and Cross, 1954 vs Kuhajda et al., 2007). As a result, the taxonomic status of Pallid and Shovelnose Sturgeon is contentious. In this study, we delineate Pallid Sturgeon and Shovelnose Sturgeon on morphological and meristic criteria exclusively, which is consistent with the typological species concept (Mayr 1996), since these can be determined immediately and objectively in the field with live specimens, and in accordance with a methodology in our use and by the same personnel since 1997 (Murphy et al., 2007). Although species determinations were made morphologically, tissue samples were collected and archived for future genetic analysis.

A numbered Floy t-bar anchor tag with a toll-free phone number was inserted externally behind the dorsal fin of all sturgeon collected. Pallid Sturgeon were electronically scanned for the presence of a Coded Wire Tag to determine if individuals were of hatchery origin from the Missouri River basin and an Avid Passive Integrated Transponder (PIT) tag indicating recapture. If no tags were detected, a non-encrypted PIT tag was inserted at the base of the dorsal fin. All sturgeon were transported in an aerated live well from the BCS and released alive back into the Mississippi River. Dead sturgeon recovered in the BCS were recorded and all Pallid Sturgeon were preserved and archived at the Engineer Research and Development Center in Vicksburg, MS or Mississippi Museum of Natural Science in Jackson, MS.

### Sturgeon Movements within the floodway

Acoustic telemetry was used to monitor movement of Shovelnose Sturgeon entrained during the 2011 opening from summer 2011 to summer 2012. Twelve VEMCO VR2Ws remote receivers were deployed into the BCS down Barbars Canal to Lake Pontchartrain to establish an automated acoustic telemetry array. Eighteen Shovelnose Sturgeon ranging in size from 501-830 mm FL were captured from upper Barbars, Y-Canal, and the BCS stilling basin and equipped with acoustic telemetry tags (V9 coded acoustic transmitters, 289 day battery life) during the period 20-27 June 2011. Tagged fish were then redistributed within the system near telemetry buoys (Barbars 1, 2, 4, 5, 8 and Y-Canal 1, see Figure 4).

### Water quality

After closure, water temperature (°C), dissolved oxygen (mg/l), and turbidity (NTU) were measured at each sampling site including the canals, Stilling Basin, and lakes with a YSI Pro DSS. In addition, water quality parameters were assessed in the Stilling Basin routinely following the closure of the BCS structure in 2018 through 2019 using a YSI Pro DSS. Point measurements were generally taken daily during early morning hours (0700-0900) from the gantry crane at bays 45, 132 and 306. These measurements were taken to better assess water quality changes, particularly dissolved oxygen, in the Stilling Basin after low bays stopped leaking to better predict and prioritize the timing of future rescue operations before conditions worsened.

### Associations between catch and structure operation

Three variables of the flood regime were calculated for each opening of the structure: onset (water temperature) in the Mississippi River, duration (days structure was open), and magnitude (volume of water passed through the structure). Water temperature of the Mississippi River during each opening was obtain from the USGS gage at Baton Rouge, LA (07374000). Duration and magnitude was obtained from the U.S. Army Corps of Engineers New Orleans District. Magnitude was the cumulative value of number of bays open converted to billion cubic meters of water. Bivariate plots between the three flood regime variables and sturgeon catch were constructed and linear regression models were calculated. Due to the limited number of observations, we use these results as exploratory and descriptive tools and not as a statistically rigorous means of testing hypotheses.

## Results

### Number of sturgeon collected

A total of 70 days with crew number ranging from 6 to 12 were expended to rescue sturgeon during the five operations (Table 2). Number of days expended per opening ranged from 5 in 2018 to 25 in 2008. Once the high bays stopped leaking allowing access to the canals and Stilling Basin, overall effort was dependent on the number of days low bays leaked after closure. Leakage provided rheophilic cues for upstream movement of sturgeon and maintained normoxic conditions, but once leakage stopped, water warmed and became hypoxic. Greater effort occurred in 2008 due to 32 days of leakage compared to 6-9 days in the four other years (Table 2). However, longer sampling days in 2008 was partly due to the development of novel sampling strategies to maximize catch, which were later refined during subsequent openings.

**Table 2.** Sampling effort by gear expended each year to capture sturgeon after operation of the Bonnet Carré Spillway. Number of days leaking was confined to the low bays once the structure was closed until the last sturgeon was captured. The 2019 values represent the second opening-closing because access was restricted after the first opening-closing.

| Year | Electroshocking,<br>hours | Gill/Trammel<br>Nets, hours | Conventional<br>Seine, hauls | Gillnet<br>Seine,<br>hours | Number of<br>Days<br>Sampled | Number<br>of<br>Leaking<br>Days |
| --- | --- | --- | --- | --- | --- | --- |
| 2008 | 15.0 | 160 | 35 | 2 | 25 | 32 |
| 2011 | 8.2 | 30 | 10 | 16.5 | 12 | 7 |
| 2016 | 5.5 | 167 | 11 | 12 | 18 | 9 |
| 2018 | 2.7 | 7 | 0 | 8 | 5 | 7 |
| 2019 | 5.6 | 8 | 10 | 25 | 10 | 6 |
| Total | 37.0 | 372 | 66 | 63.5 | 70 | 61 |

Electroshocking was the most effective gear to catch sturgeon in the canals and lakes, ranging from hours in 2018 to 15 hours in 2008. Thirty-six percent of the Pallid Sturgeon were collected by electroshocking. Gillnets and trammel nets were set in the canals and lakes for a total of 372 hours over the five openings collecting 22% of the Pallid Sturgeon. The modified gillnet-seine was pulled for a total of 63.5 hours primarily in the Stilling Basin collecting 15% of the Pallid Sturgeon. Castnets and dipnets collected 13% and 11% of the Pallid Sturgeon, respectively. Other techniques accounted for less than 5%. Gear efficiency was similar for Shovelnose Sturgeon.

A total of 57 Pallid Sturgeon and 362 Shovelnose Sturgeon were collected after the five operations (Table 1). Fork length (mm) ranged from 528-1038 and 384-928 for Pallid and Shovelnose Sturgeons, respectively. A notable collection was a tagged Pallid Sturgeon originally captured in the floodway during 2008, released back into the Mississippi River, and recaptured in the floodway in 2011. Number of Pallid Sturgeon collected ranged from zero in 2016 after the winter opening to 20 in 2011. Shovelnose Sturgeon were also not collected in 2016 but 219 individuals were collected in 2019, almost 3 times the number collected in other years.

### Sturgeon movements within the floodway

The telemetry array was deployed from 20 June 2011 through 25 August 2012 and accumulated over 120,000 detections. No mortalities were observed following the tagging period and initially all individuals moved extensively near their original release point. The initial acoustic array (n = 10 receivers) within the floodway was deployed on 20 June 2011 prior to sampling. The remaining receivers near Lake Pontchartrain, two new receivers and one receiver relocated from the upper floodway, were not deployed until 13 July. This created an “open window” for undocumented movement into Lake Pontchartrain with 6 individuals unaccounted for after 13 July suggesting they moved quickly through the floodway and into Lake Pontchartrain before the final receivers were deployed. None were documented returning to the floodway. Alternatively, the lack of detections during this period could be due to tag failure. Those fish that remained in the system experienced sporadic, localized movement with no detection patterns to support movement of telemetry tagged individuals from the BCS into Lake Pontchartrain after 13 July. However, overall movement of telemetry tagged fish began to decrease by early August, as water levels within the floodway decreased, in part creating isolated pools and remnant channels, and as water temperatures increased (31° C). This pattern of decreased movement was also likely in response to the loss of rheophilic cues as bay leakage at the BCS was minimal to none resulting in decreased water flow through the entire floodway. Salinity during this period where the floodway enters Lake Pontchartrain was ≥ 2 ppt; detections during this period on the receivers nearest to Lake Pontchartrain were few to none.

### Water quality

The structure was operated in late winter (2016 and first opening of 2019), spring only (2008, 2011, and 2018), and spring into mid-summer (second opening of 2019). As a result, mean water temperature in the floodway varied from 10.8 C during the winter operation of 2016 to a high of 30.8 C in 2019 when the operation extended into the summer. However, mean water temperature in the Mississippi River was consistently lower when the structure was open compared to measurements taken after closure (Table 1). Temperature difference was most pronounced in 2018 when the floodway was 11°C higher compared to the river. Although mean dissolved oxygen in the floodway was normoxic during all years, diel fluctuations did occur in the stilling basin and lakes resulting in hypoxic (< 3 mg/l) conditions.

Leakage through the bays during closure occurs most years, ranging from near zero in 2012 and 2014 to 60% during 2019, affecting water quality in the stilling basin and canals (Figure 5). Leakage of Mississippi River water into the BCS moderates temperatures and prevents hypoxia. Water quality monitoring in the stilling basin during 2018 and 2019 clearly showed a rapid decrease in dissolved oxygen when the low bays quit leaking (Figure 6). Once the low bays quit leaking in August, the stilling basin became hypoxic during part of the day creating physiological stress on trapped sturgeon and other fish species leading to fish kills. Mississippi River stage elevations are monitored to determine when leaking through the low bays will end ensuring that rescue operations in the stilling basin are scheduled accordingly.

**Figure 5.**
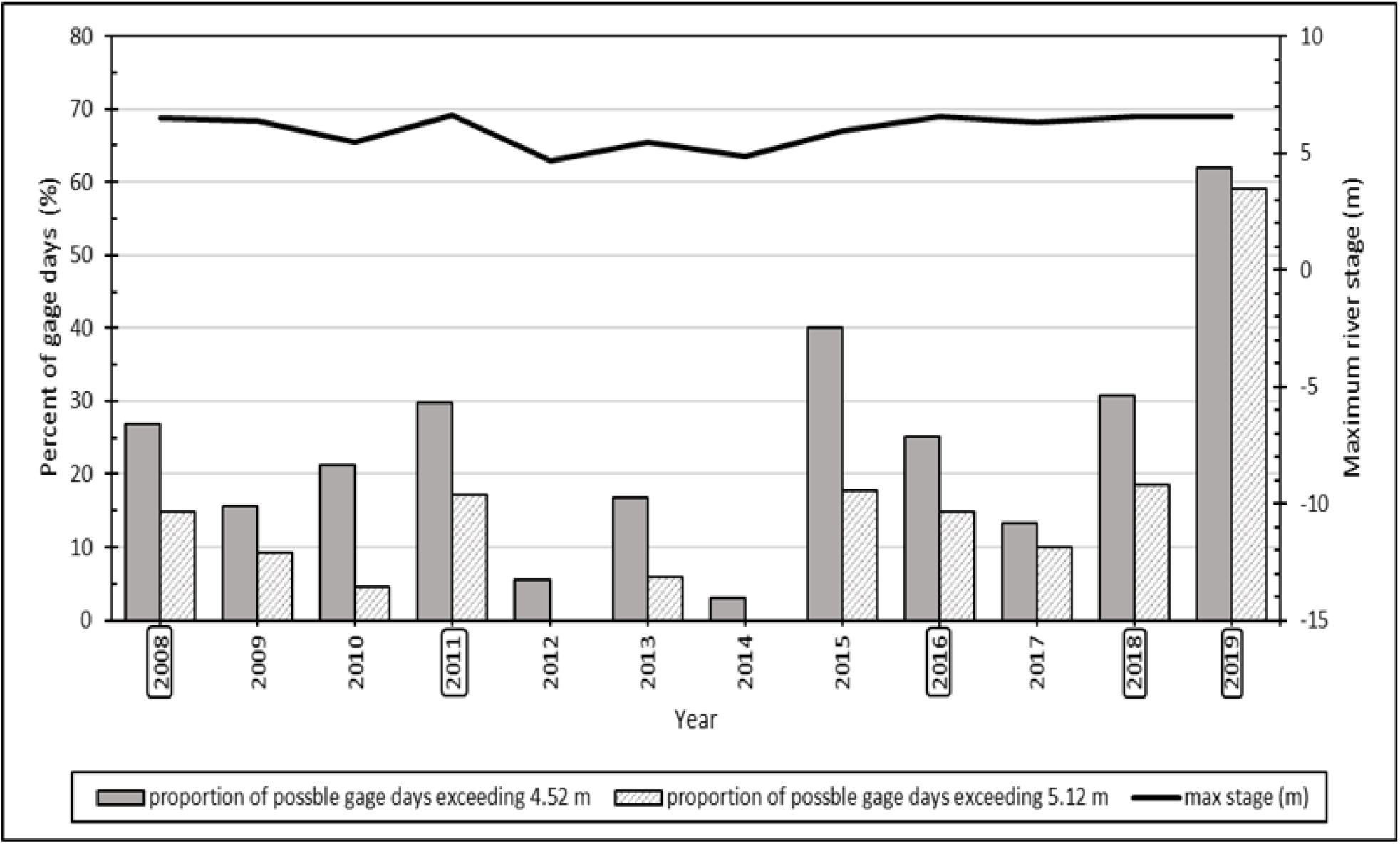
Percentage of gage days each year from 2008 – 2019 that water leaked through the pins at the low bays (gage > 4.52 m) and high bays (gage >5.12 m). Some years had incomplete gage records and therefore percent gage days during each year based on available records was used to denote leakage. Mississippi River stage shown for the Bonnet Carré gage (01280). Years when the Bonnet Carré structure was open are outlined.

**Figure 6.**
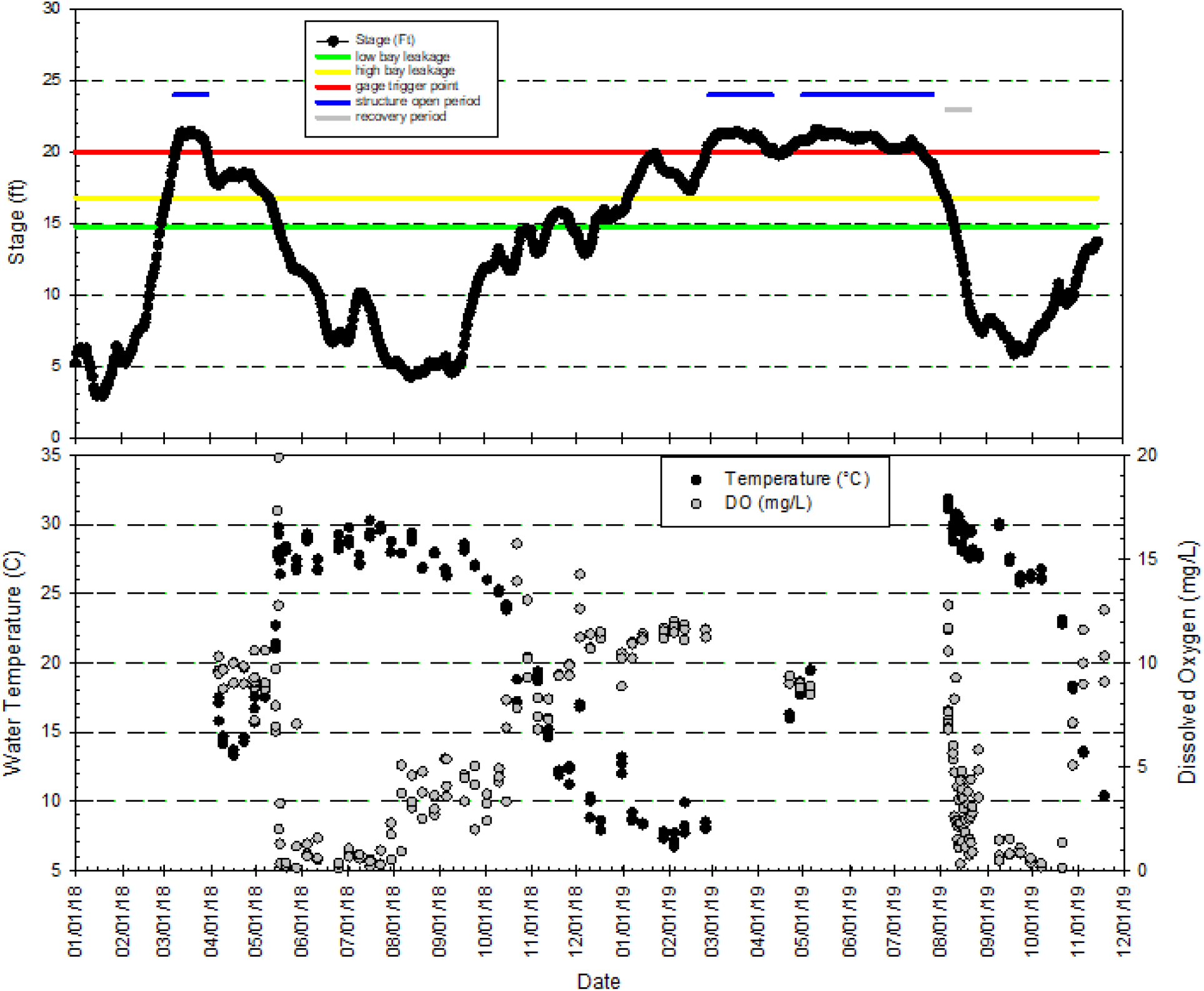
Pattern of leakage (top panel) and water temperature and dissolved oxygen measurements (bottom panel) taken in Bonnet Carre’ Spillway Stilling Basin from April 2018 through November 2019. Continuous leakage through the low bays was noted from 14 December 2018 through 10 August 2019.

### Associations between sturgeon catch and structure operations

Numbers of Pallid and Shovelnose Sturgeon collected were positively correlated with onset (i.e., date of initial opening), duration, and magnitude of Bonnet Carré openings with coefficients of determination ranging from R^2^=0.58 to R^2^=0.99. (Figure 7). For Pallid Sturgeon, correlation was highest for onset. Entrainment risk is negligible at water temperatures below 10 °C and high as water temperatures increase above 20 °C. The duration and magnitude of openings were more curvilinear, suggesting that number of individual Pallid Sturgeon tended to plateau after prolonged openings. For Shovelnose Sturgeon, correlation was highest for duration, indicating that depletion of the riverine population was less likely for the more abundant Shovelnose Sturgeon. The variation in catch of Shovelnose Sturgeon between 2019 and 2011 as a function of water volume passing through the structure also indicates that number entrained may continue to increase as magnitude increases.

**Figure 7.**
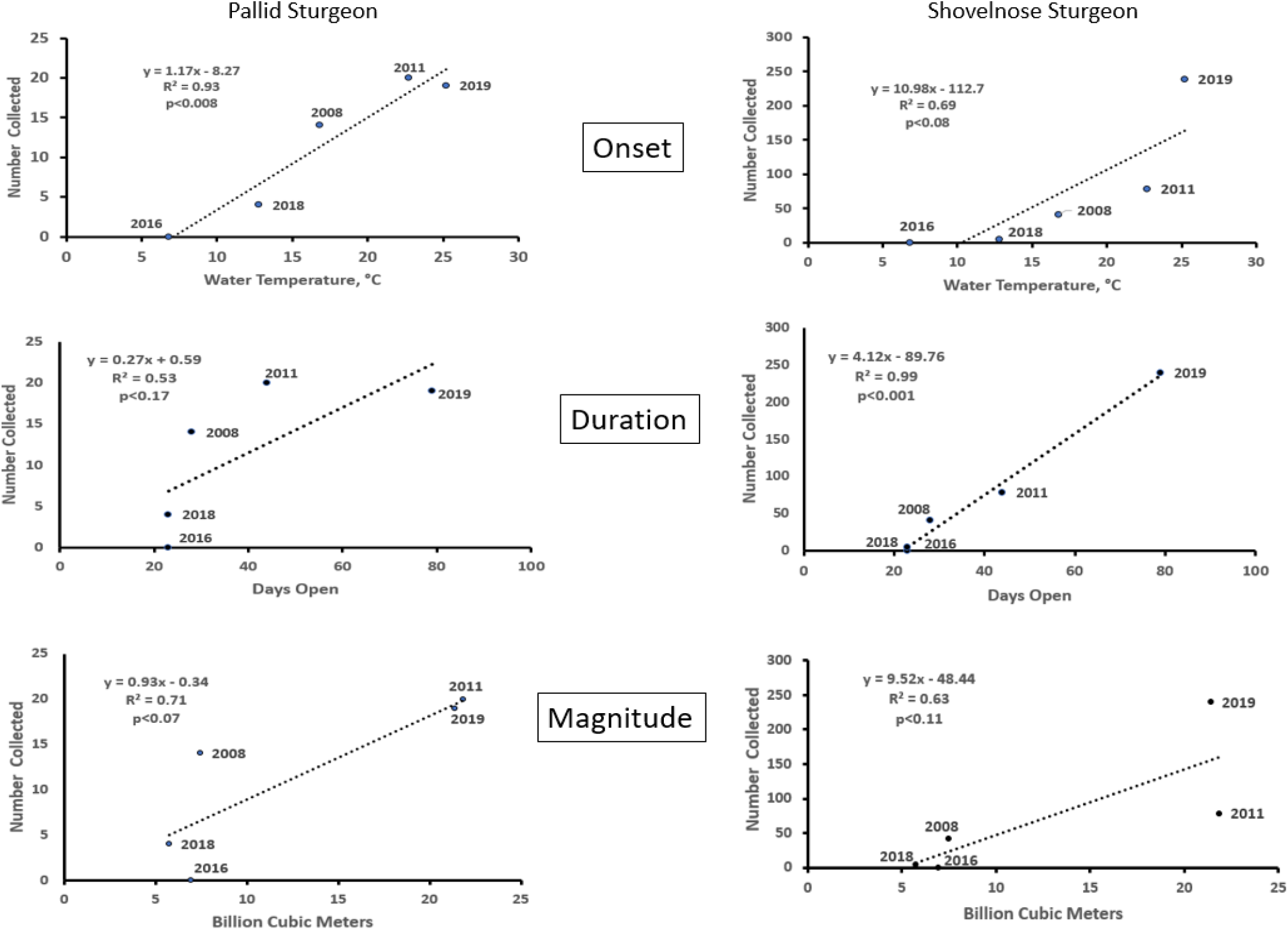
Regression analysis of number of Pallid Sturgeon and Shovelnose Sturgeon captured as a function of Mississippi River mean water temperature (°C) at Baton Rogue, LA during operation of the Bonnet Carré Spillway (Onset), number of days the structure was open (Duration), and total volume (billion cubic meters) of water passed through the structure (Magnitude). Year indicated by each data point.

## Discussion

Rescuing Pallid Sturgeon from waterbodies in the BCS required perseverance under constantly changing conditions even on a daily basis. The five openings of the BCS had different operating and environmental conditions, and as a result, different outcomes in the number of sturgeon collected. A major consideration for sampling was the amount and duration of leakage through the bays that maintains discharge in the canals. Bay leakage depends on Mississippi River stage elevation relative to the crest of the high and low bays. Pallid and Shovelnose Sturgeon are inherently strongly rheotactic (Adams et al. 1998, 1999; Parsons et al., 2003; Hoover et al. 2011). Consequently, they will orient in the direction of the flow towards the structure. As flow declines and bay leakage diminishes, canals and shallow lakes become isolated due to accretion of sediment plugs during the opening of the structure creating potential barriers to upstream passage. Without flow, sturgeon may become trapped in hypoxic lakes and canals in the floodway or wander into Lake Pontchartrain where they likely perished due to the inability to osmoregulate in saline waters. Longer flow duration in the canals will lead to higher capture rates.

No sturgeon were captured after the 2016 openings when water temperatures were much colder and apparently fish were not moving along the channel border of the Mississippi River where they are more susceptible to entrainment. Pallid and Shovelnose Sturgeon catch rates in the Mississippi River on trotlines generally increase as water temperature approaches 10 °C (Killgore et al. 2007). The mean water temperature of the Mississippi River during January and February 2016 when the spillway was open ranged from 7.5 to 7.8 °C (USGS gage 07374000, Baton Rouge, LA). Although mean water temperature in the spillway during rescue operations was 10.8 °C (Table 1), colder temperatures in the river persisted during the opening period when sturgeon are more inactive. Higher temperatures during other operations resulted in more sturgeon entrained, particularly Shovelnose Sturgeon that were caught more frequently than Pallid Sturgeon when temperatures rise above 20 C (Killgore et al. 2007). Pallid Sturgeon occupy the main channel primarily during low river stage and warm temperature conditions that occur in summer and early autumn according to a telemetry study in the Lower Mississippi River (Herrala et al. 2014). At higher river stages, both species may be more inclined to move along the channel border closer to the spillway regardless of water temperature.

Sturgeon approaching the open spillway encounter entrainment velocities as water overtops the concrete weir, which can exceed 2 m/s (USACE New Orleans District, personnel communication). Adult Shovelnose Sturgeon and presumably Pallid Sturgeon exploit boundary-layers along the substrate to effectively move or hold position in fast-flowing rivers. Both species, in laboratory studies, show relatively weak prolonged swimming ability. Adult Shovelnose Sturgeon have 60-minute and 15-minute critical swimming speeds of only 0.6 m/s and 0.6-1.2 m/s in open water, respectively, and 1.3-1.7 m/s in boundary layers (Parsons et al., 2003; Hoover et al., 2011; Adams et al., 1997). Juveniles of both species have 30-min critical swim speeds < 0.4 m/s (Adams et al., 1999; Adams et al, 2003). Relative weak swimming capability in the vicinity of fast entrainment velocities render these species vulnerable to entrainment anywhere near the structure. Differences in swimming behavior between the two species have been noted, however, which may increase vulnerability to entrainment. Shovelnose Sturgeon tend to free-swim in the water column when reaching higher swimming speeds compared to Pallid Sturgeon that hunker down (Adams et al 2003). Free-swimming would increase risk of entrainment compared to skimming along the substrate or station holding.

Nineteen Pallid Sturgeon were collected in 2019, similar to 2011, whereas 239 Shovelnose Sturgeon were collected in 2019 almost 3 times higher than 2011 when 78 Shovelnose Sturgeon were collected. The Pallid to Shovelnose ratio in the lowermost reach of the lower Mississippi River is typically 1:3 (Killgore et al. 2007) but was 1:12.6 in 2019 (Table 1). Although the 2019 openings from February to April and May to July may have coincided with one or more major movements and dispersal of Shovelnose Sturgeon, depletion of Pallid Sturgeon in the vicinity of the structure is also a consideration. Number of Pallid Sturgeon collected was similar in 2011 and 2019 when the highest water volumes passed through the structure suggesting a depletion of individuals that could be entrained. Conversely, number of Shovelnose Sturgeon collected was variable at higher water volumes suggesting that numbers entrained may continue to increase at even higher volumes passing through the structure (Figure 7).

The stilling basin becomes a hypoxic death trap for sturgeon after bays stop leaking. It is a rectangular concrete structure 2,347 m long, 9.1 m wide, averaging 1.2 m in depth, and holds water year around. There is no escape for stranded fish unless bays begin leaking again to a point that reconnects the downstream canals as Mississippi River stages rise. Of the 57 Pallid Sturgeon collected during the five openings, four were dead with one found in the stilling basin. Of the 362 Shovelnose Sturgeon collected, 98 were found dead, and of these, 92% were collected in the stilling basin. Fish kills typically occur in the stilling basin after each operation as water temperature rises and dissolved oxygen decreases but are often species-specific. For example, tens of thousands of Skipjack Herring (*Alosa chrysochloris*) became stranded in the stilling basin and died in 2018. Sturgeon are also sensitive to low dissolved oxygen and hypoxic conditions impair their respiratory metabolism, foraging activity, and growth rates (Cech and Doroshov 2004). Blevins (2011) reported that recruitment of Pallid Sturgeon in the Missouri River may be limited by high summer water temperatures in excess of 30 °C and dissolved oxygen concentrations less than 2 mg/l in late spring and summer. Therefore, rescue operations in the stilling basin must begin before water becomes hot and hypoxic to minimize sturgeon mortality (Figure 8).

**Figure 8.**
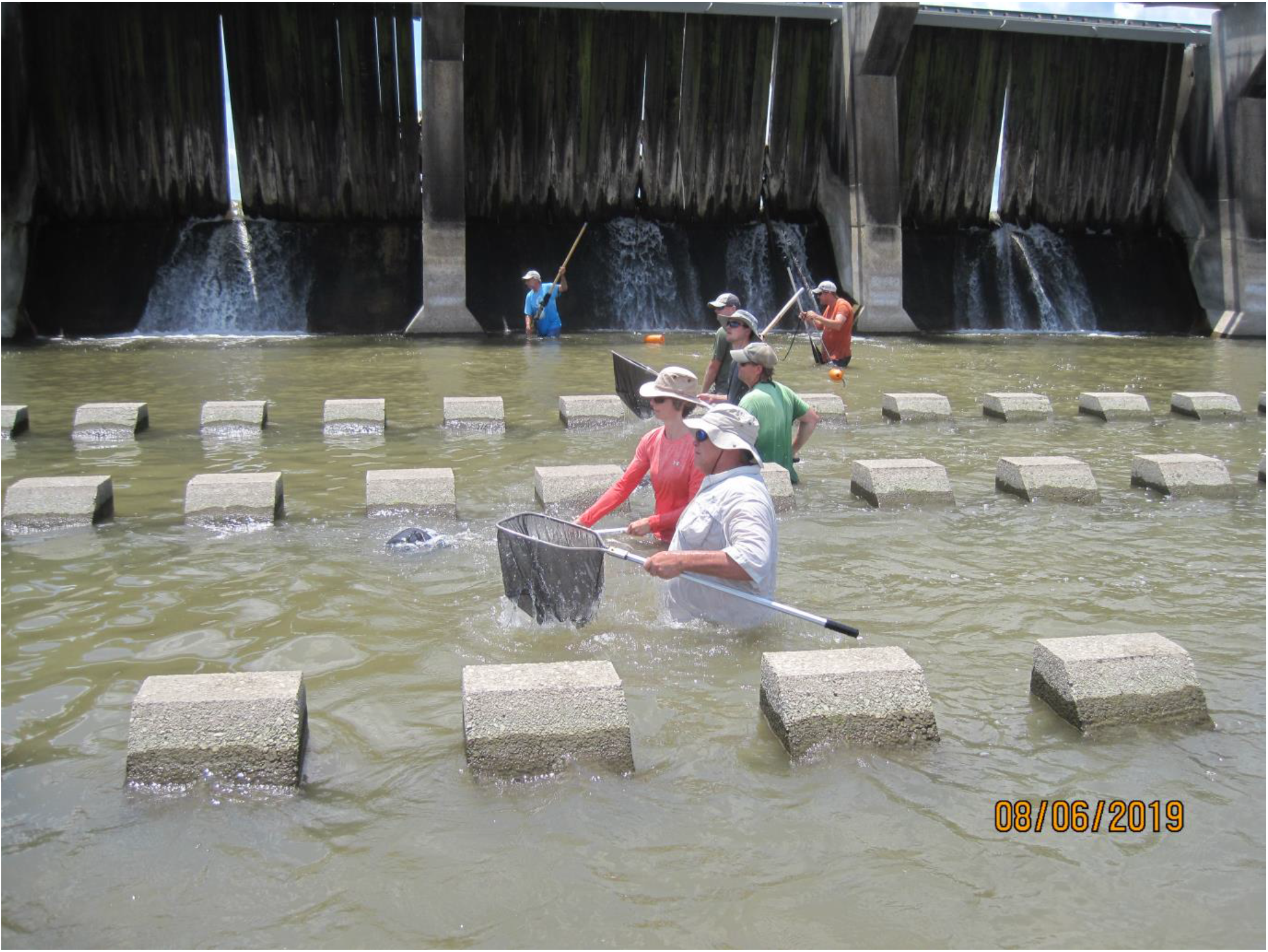
Collecting efforts in the stilling basin require at least 10 people - 3 pulling the modified gillnet-seine in the wide section nearest the structure, four wading between the baffles with large dipnets, one data recorder, and a transport team assisting with sturgeon measurements, tagging, and release back into the Mississippi River. Two additional people moving a block net to corral fish will usually increase capture rates. Leakage from the Mississippi River thru pins is evident.

Entrainment risk is related to duration and seasonality of openings. Capture rate is initially high but over time declines. Several reasons could explain this trend. Sampling efficiency and effort varies among collection periods. However, the most likely explanations are that sturgeon are displaced further downstream in the floodway, become trapped in isolated waterbodies, lack rheophilic cues as flow subsides, or wander into Lake Pontchartrain during longer openings and cannot find their way back into the spillway where they are more likely to be rescued. This suggests that the number captured in the spillway after closure is an underestimate of the total number entrained. A conservative estimate of Pallid Sturgeon age 3+ population size in the 1,931-km reach of the Mississippi River below the confluence of the Missouri River ranged from 4.5–15 fish per river kilometer or a total of 4,600 to 15,000 (Friedenberg et al. 2017). Hintz et al. (2016) estimated population size of Pallid and Shovelnose Sturgeon in the Middle Mississippi River, a 322-km reach between the confluences of the Missouri and Ohio Rivers, at 1,516 (5 individuals/rkm) and 82,336 (266 individuals/rkm), respectively. The annual population estimate for wild Pallid Sturgeon in an 877-km reach of the Lower Missouri River varied from 5.4 to 8.9 fish/rkm, whereas the estimate for known hatchery-reared fish varied from 28.6 to 32.3 fish/rkm (Steffensen et al. 2012). The relatively small number of Pallid and Shovelnose Sturgeon rescued from the BCS represent less than 1% of the total population size in the Lower Mississippi River with even the lowest estimates. However, adding hundreds that may have been entrained but not captured could lead to impacts on overall population size if the BCS continues to be opened on a more frequent basis.

Under the authority of the Federal Endangered Species Act, the USFWS has issued several Section 7 “No jeopardy biological opinions” on opening the Bonnet Carre’, which essentially means that entrainment of Pallid and Shovelnose Sturgeon through the structure is not likely to jeopardize their continued existence. However, Section 7(a)(1) of the Act directs Federal agencies to “utilize their authorities to further the purposes of the Act by carrying out conservation programs for the benefit of endangered and threatened species.” Conservation recommendations are discretionary agency activities to minimize or avoid adverse effects of a proposed action on listed species or critical habitat, to help implement recovery plans, or to develop information. Based on this study, recommended conservation measures include:

a. Continue rescue operations after each operation using the collecting made in this article, methods and timing made in this article,
b. Maintain or construct a defined channel (s) in the upper reach of the floodway with adequate depths to enhance directional cues for sturgeon moving back towards the structure and to provide long-term navigability for collection vessels once the structure is closed,
c. Use pumps and/or siphons to transfer river water and circulate the Stilling Basin when Mississippi River water levels drop below the concrete weir on the structure to improve water quality conditions thereby reducing stress on entrained sturgeon prior to rescue, and,
d. Utilize acoustic telemetry to evaluate movement rates and patterns in the floodway, as well as dispersal potential into Lake Pontchartrain. Acoustic tags can be implanted in Shovelnose Sturgeon as surrogates for Pallid Sturgeon.

There is no doubt that the BCS will operate again as flood frequencies increase in the Lower Mississippi River. Multiple observations have confirmed that the capture of entrained sturgeon and other chondrostean fish in the floodway and release back into the Mississippi River is a viable solution to reduce population impacts. Injury from freefall from passing over a spillway causing abrasions and scrapes may affect survival (Rytwinski et al. 2017). However, the recapture of a Pallid Sturgeon in 2011 (this study) indicates annual survival of rescued individuals and all sturgeon released back into the Mississippi River swam away under their own volition. Furthermore, an adult Paddlefish (*Polydon spathula*) entrained through the structure in 2011, which was injured and underweight, and recaptured eight months later in northern Mississippi near Greenville, 627 km upriver from where it was released, indicates that a large entrained fish, trapped for several days in a hyperthermic and hypoxic habitat, can be viable when returned to the river (Hoover et al., 2013). The experience gained over the past 5 operations will ensure that rescue operations will continue in an effective manner and compliance with the Endangered Species Act will be one of the priorities in fighting floods on the Lower Mississippi River.

## Acknowledgments

Logistical requirements of this study required a large number of agencies and individuals assisting with sturgeon rescue operations and permitting: U. S. Army Corps of Engineers New Orleans District / Bonnet Carre Staff - Richard Boe, Michael Brown, Tony Catalanotto, Richard Cusimano, Rob Heffner, Emile "Skip" Jacobs, Tim Lacoste, Howard Ladner, Bill Maus, John “Rusty“ Munson, Thomas Parker, Michael Saucier, and Steve Stone; Louisiana Department of Wildlife and Fisheries - Matt Duplessis, Robby Maxwell, Patrick Morris, Alex Perret, Tim Ruth, Brac Salyers, Jeff Thompson, Gary Vitrano, and Jonathan Winslow; U.S. Fish and Wildlife Service - Debbie Fuller, Paul Hartfield, Monica Sikes, Rob Smith, Karen Soileau, David Walther; Nicholls State University - Stephen Byrne, Chris Levron, Dave Shultz, and Clint Troxler; U.S. Army Engineer Research and Development Center - Krista Boysen, Jay Collins, Nicky Faucheux, Chris Giesler, Alan Katzenmeyer, Phil Kirk, Bill Lancaster, Bradley Lewis, Catherine Murphy, Amanda Oliver, and Max Wamsley. Funding was provided by the U. S. Army Corps of Engineers New Orleans District and the Mississippi River Geomorphology and Potamology Program at the U. S. Army Mississippi Valley Division.

